# APC/C^FZR-1^ Controls ZYG-1 Levels to Regulate Centrosome Assembly

**DOI:** 10.1101/2020.08.12.248658

**Authors:** Jeffrey C. Medley, Joseph R. DiPanni, Luke Schira, Blake M. Shaffou, Brandon M. Sebou, Mi Hye Song

## Abstract

Aberrant centrosome numbers are associated with human cancers. The levels of centrosome regulators positively correlate with centrosome number. Thus, tight control of centrosome protein levels is critical. In *Caenorhabditis elegans*, the anaphase-promoting complex/cyclosome and co-activator FZR-1 (APC/C^FZR-1^) ubiquitin ligase negatively regulates centrosome assembly through SAS-5 degradation. In this study, we identify the *C. elegans* ZYG-1 (Plk4 in human) as a new substrate of APC/C^FZR-1^. Inhibiting APC/C^FZR-1^ or mutating a ZYG-1 destruction (D)-box leads to elevated ZYG-1 levels at centrosomes, restoring bipolar spindles and embryonic viability to *zyg-1* mutants, suggesting that APC/C^FZR-1^ targets ZYG-1 for proteasomal degradation via D-box motif. We also show the Slimb/βTrCP-binding (SB) motif is critical for ZYG-1 degradation, substantiating a conserved mechanism by which ZYG-1/Plk4 stability is regulated by SCF^Slimb/βTrCP^-dependent proteolysis via the conserved SB motif in *C. elegans*. Furthermore, inhibiting both APC/C^FZR-1^ and SCF^Slimb/βTrCP^, by co-mutating ZYG-1 SB and D-box motifs, stabilizes ZYG-1 in an additive manner, conveying that APC/C^FZR-1^ and SCF^Slimb/βTrCP^ ubiquitin ligases function cooperatively for timely ZYG-1 destruction in *C. elegans* embryos where ZYG-1 activity remains at threshold level to ensure normal centrosome number.

## Introduction

Centrosomes, as the primary microtubule-organizing centers, establish bipolar mitotic spindles that ensure accurate transmission of genomic content into two daughter cells. To sustain genomic integrity, centrosome number must be strictly controlled by duplicating only once per cell cycle. Abundance of centrosome factors directly influences centrosome number. Blocking degradation of centrosome factors causes extra centrosomes, while their depletion inhibits centrosome duplication (Nigg and Holland, 2018). The ubiquitin-proteasome system provides a key mechanism to control centrosome protein levels (Nakayama and Nakayama, 2006). Levels of centrosome factors are regulated by proteasomal degradation through E3 ubiquitin ligases, including the Anaphase Promoting Complex/Cyclosome (APC/C) and SKP1-CUL1-F-box-protein (SCF), and their regulatory mechanisms appear to be conserved (Arquint et al., 2012; Cunha-Ferreira et al., 2009; Holland et al., 2010; Medley et al., 2017a; Meghini et al., 2016; Peel et al., 2012; Puklowski et al., 2011; Rogers et al., 2009; Strnad et al., 2007; Tang et al., 2009). In *C. elegans*, the kinase ZYG-1 is a key centrosome regulator and ZYG-1 levels are critical for normal centrosome number and function (O’Connell et al., 2001; Song et al., 2008). Abundance of Plk4 (human homolog of ZYG-1) is regulated by the SCF-mediated proteolysis through the F-box protein Slimb/βTrCP (*Drosophila/humans*) (Cunha-Ferreira et al., 2009; Holland et al., 2010; Rogers et al., 2009). Such mechanism is conserved in *C. elegans* where ZYG-1 levels are controlled by SCF^Slimb/βTrCP^ (Peel et al., 2012).

Another E3 ubiquitin ligase, APC/C, also regulates levels of centrosome regulators. In *C. elegans*, FZR-1 (Cdh1 in human), a coactivator of APC/C, negatively influences centrosome assembly, and APC/C^FZR-1^ regulates SAS-5 levels through KEN-box (Kemp et al., 2007; Medley et al., 2017a), which is conserved in humans and flies where APC/C^Cdh1/Fzr^ targets centrosome factors (human STIL/SAS-5, SAS-6, CPAP/SAS-4 and *Drosophila* Spd2) via KEN-box (Arquint and Nigg, 2014; Meghini et al., 2016; Strnad et al., 2007; Tang et al., 2009). Here, we investigate ZYG-1 as a potential APC/C^FZR-1^ substrate, and how E3 ubiquitin ligases APC/C^FZR-1^ and SCF^Slimb/βTrCP^ influence ZYG-1 levels in centrosome assembly.

## Results and Discussion

### Loss of FZR-1 leads to Elevated Centrosomal ZYG-1 Levels

Prior study identified SAS-5 as an APC/C^FZR-1^ substrate, and implicated that APC/C^FZR-1^ might target additional centrosome regulators in *C. elegans* (Medley et al., 2017a). FZR-1 was identified as a genetic suppressor of ZYG-1 since loss of FZR-1 restores embryonic viability and centrosome duplication to *zyg-1(it25)* mutants (Kemp et al., 2007; Medley et al., 2017a). Given the strong genetic interaction between *zyg-1* and *fzr-1*, a compelling possibility is that ZYG-1 might be an APC/C^FZR-1^ substrate. In *C. elegans* embryos, while cellular ZYG-1 protein remains low, centrosome-associated ZYG-1 is regulated during cell cycle, peaking at anaphase and declining toward mitotic exit (Song et al., 2008, 2011). ZYG-1 levels are shown to be regulated by protein phosphorylation (Song et al., 2011) and proteasomal degradation (Peel et al., 2012).

If ZYG-1 is degraded by APC/C^FZR-1^, downregulation of APC/C^FZR-1^ should protect ZYG-1 from destruction leading to protein accumulation. To address this, we stained embryos with anti-ZYG-1 and quantified fluorescence intensity of centrosome-associated ZYG-1 signal (**Fig. 1**). Quantitative immunofluorescence (IF) reveals that, compared to wild-type (WT) controls (1.00 ± 0.25 fold), *fzr-1* mutant embryos exhibit increased levels of centrosomal ZYG-1 during the first anaphase (1.28 ± 0.27 fold). *zyg-1(it25)* contains a temperature-sensitive point mutation (P442L) within the cryptic polo-box (CPB) domain that is critical for centrosomal localization of ZYG-1 (Fig 2A) (Kemphues et al., 1988; O’Connell et al., 2001; Shimanovskaya et al., 2014). In *zyg-1(it25)* embryos, centrosomal ZYG-1 is reduced to ~40% of WT controls. By contrast, centrosomal ZYG-1 levels in *zyg-1(it25) fzr-1(bs31)* double-mutants are significantly increased (0.80 ± 0.33 fold) compared to *zyg-1(it25)* mutants, indicating the *fzr-1* mutation leads to partial restoration of centrosomal ZYG-1 to *zyg-1(it25)* embryos. Furthermore, inhibition of MAT-3/APC8, an essential subunit of the APC/C complex (Golden et al., 2000) results in elevated centrosomal ZYG-1, supporting that FZR-1 regulates centrosomal ZYG-1 through an APC/C-dependent mechanism (**Fig. S1A**).

**Figure 1.**
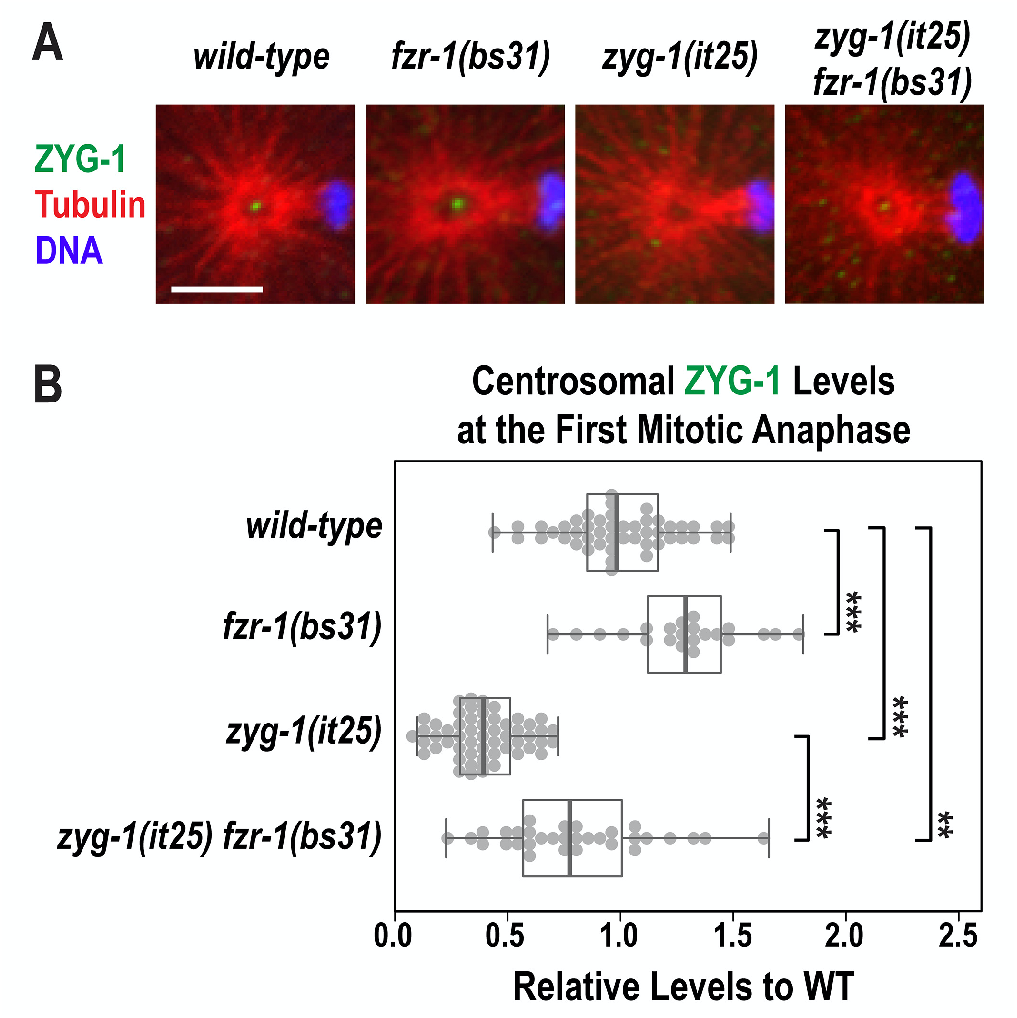
Loss of FZR-1 results in elevated centrosomal ZYG-1 levels. (A) Anterior centrosomes stained for ZYG-1 at the first mitotic anaphase. Bar, 5 μm. (B) Quantification of centrosomal ZYG-1 levels. In the plot, box ranges from the first through third quartile of the data. Thick bar indicates the median. Solid grey line extends 1.5 times the inter-quartile range or to the minimum and maximum data point. Each dot represents a centrosome. \*\**p*<0.01, \*\*\**p*<0.001 (two-tailed t-tests).

**Figure 2.**
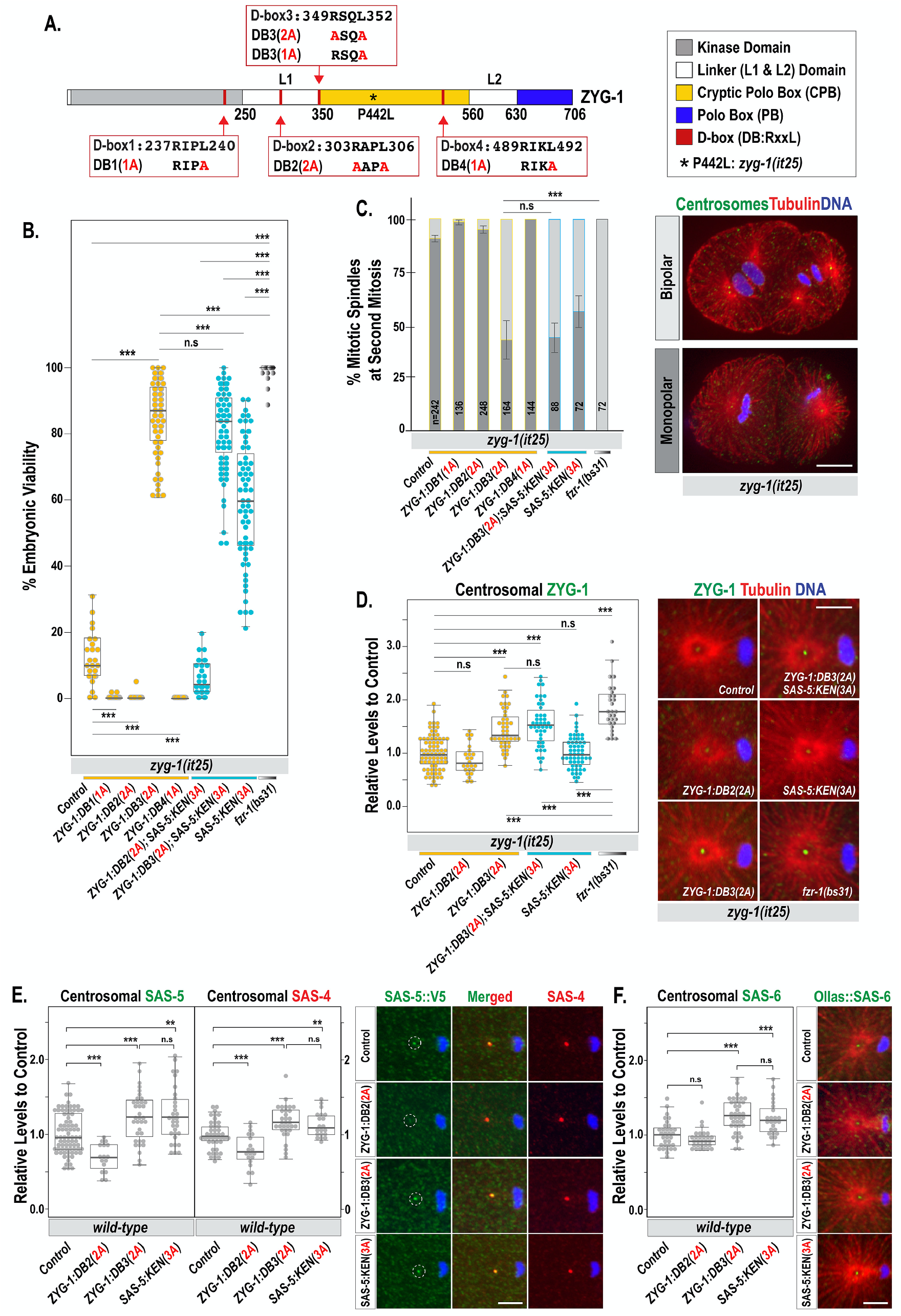
Mutating ZYG-1 D-box3 leads to the *zyg-1* suppression and elevated centrosomal ZYG-1 levels. (A) ZYG-1 protein structure illustrates the locations of D-boxes and alanine-substitutions, and functional domains. (B) Embryonic viability at 22.5°C (Table 1). Each dot represents a hermaphrodite. (C) Quantification of monopolar (dark grey) and bipolar (light grey) spindles in two-cell embryos at 23°C. Average and standard deviation (SD) are presented. n is the number of blastomeres (left). Embryos stained for centrosomes (ZYG-1), microtubules and DNA, illustrate mitotic spindles at second division. Bar, 10 μm (right). (D) Quantification of centrosome-associated ZYG-1 levels at the first mitotic anaphase (left). Centrosomes stained for ZYG-1 at the first anaphase. Bar, 5 μm (right). (E,F) Quantification of centrosomal levels of SAS-5 (E), SAS-4 (E), and SAS-6 (F) at the first mitotic anaphase. (D-F) Each dot represents a centrosome; Shown are anterior centrosomes. Bar, 5 μm. (B,D-F) In the plots, box ranges from the first through third quartile of the data. Thick bar indicates the median. Solid grey line extends 1.5 times the inter-quartile range or to the minimum and maximum data point. ^ns^*p*>0.05, \*\**p*<0.01, \*\*\**p*<0.001 (two-tailed t-tests).

These results support a hypothesis that ZYG-1 levels are regulated by APC/C^FZR-1^-dependent proteolysis. Despite exhaustive efforts to address how loss of FZR-1 impacted cellular ZYG-1 levels, we were unable to detect endogenous ZYG-1 by immunoblot or immunoprecipitation (IP) due to extremely low abundance of ZYG-1. However, IP data suggest a possible physical interaction between ZYG-1 and FZR-1 (**Fig. S1B**), consistent with a hypothesis that ZYG-1 could be a potential APC/C^FZR-1^ target.

### ZYG-1 Contains Putative Destruction (D)-box Motifs

The APC/C^FZR-1^ E3 ubiquitin ligase targets substrates via interaction of the coactivator FZR-1 with conserved motifs, predominantly the destruction (D)-box and KEN-box degrons, within the substrate (Glotzer et al., 1991; Pfleger and Kirschner, 2000). While ZYG-1 contains no KEN-box, in silico analysis identified four putative D-box motifs (RxxL) within ZYG-1. We then asked if APC/C^FZR-1^ might target ZYG-1 for degradation through D-box. In ZYG-1 (**Fig. 2A**), D-box1 resides within kinase domain, D-box2 in Linker 1 (L1) domain, D-box3 at the junction between L1 and CPB domains, and D-box4 within CPB domain (Lettman et al., 2013; O’Connell et al., 2001; Shimanovskaya et al., 2014). The sequence alignments show D-box1 is the only conserved motif in closely related nematodes (**Fig. S2**).

To determine if any of putative D-boxes might be functional degron, we mutated each D-box (RxxL) by substituting two critical residues to alanine (AxxA) at the endogenous locus in wild-type(N2) animals (**Fig. 2A**). Using CRISPR/Cas9 genome editing, we generated D-box2 and D-box3 mutants with two-alanine substitution (2A; AxxA), termed the ZYG-1^**DB2(2A)**^ and ZYG-1-^**DB3(2A)**^ mutation. However, the 2A mutation of D-box1 or D-box4 produced sterile hermaphrodites. In human cells, L45 in Cyclin B1:D-box (42RxxL45) is critical for Cyclin B1 destruction (Matsusaka et al., 2014). Thus, we generated D-box1 and D-box4 mutants with single-alanine substitution (1A; RxxA), termed the ZYG-1^**DB1(1A)**^ and ZYG-1^**DB4(1A)**^ mutation. All four ZYG-1 D-box mutations in WT background produce viable progeny (**Table 1**).

**Table 1.** Genetic Analysis.

| Strain | °C | % Embryonic Viability<br>(ave ± sd) | n<br>(Progeny) |
| --- | --- | --- | --- |
| *wild-type (N2)* | 22.5 | 99.9 ± 0.3 | 719 |
| *ZYG-1<sup>DB1(1A)</sup>* | | 99.5 ± 1.4 | 1873 |
| *ZYG-1<sup>DB2(2A)</sup>* | | 99.9 ± 0.3 | 2181 |
| *ZYG-1<sup>DB3(2A)</sup>* | | 99.8 ± 0.4 | 2174 |
| *ZYG-1<sup>DB4(1A)</sup>* | | 99.3 ± 1.7 | 1528 |
| *SAS-5<sup>KEN(3A)</sup>* | | 99.8 ± 0.6 | 2202 |
| *fzr-1(bs31)* | | 99.8 ± 0.9 | 1087 |
| *zyg-1(it25)* | | 14.0 ± 11.8 | 1953 |
| *zyg-1(it25) ZYG-1<sup>DB1(1A)</sup>* | | 0.1 ± 0.5 | 2072 |
| *zyg-1(it25) ZYG-1<sup>DB2(2A)</sup>* | | 0.1 ± 0.9 | 2042 |
| *zyg-1(it25) ZYG-1<sup>DB3(2A)</sup>* | | 85.2 ± 13.6 | 3959 |
| *zyg-1(it25) ZYG-1<sup>DB4(1A)</sup>* | | 0.0 ± 0.0 | 2295 |
| *zyg-1(it25) ZYG-1<sup>DB2(2A)</sup>; SAS-5<sup>KEN(3A)</sup>* | | 6.10 ± 5.1 | 1840 |
| *zyg-1(it25) ZYG-1<sup>DB3(2A)</sup>; SAS-5<sup>KEN(3A)</sup>* | | 81.4 ± 13.2 | 3882 |
| *zyg-1(it25); SAS-5<sup>KEN(3A)</sup>* | | 59.5 ± 18.1 | 4006 |
| *zyg-1(it25) fzr-1(bs31)* | | 99.1 ± 2.4 | 1451 |
| *zyg-1(it25)* | 22.5 | 5.4 ± 6.3 | 2325 |
| *zyg-1(it25) ZYG-1<sup>SB(S335A)</sup>* | | 16.5 ± 12.5 | 1874 |
| *zyg-1(it25) ZYG-1<sup>SB(T339A)</sup>* | | 15.4 ± 12.2 | 2177 |
| *zyg-1(it25) ZYG-1<sup>SB(2A)</sup>* | | 58.8 ± 12.1 | 2401 |
| *zyg-1(it25) ZYG-1<sup>DB3(2A)</sup>* | | 45.7 ± 9.7 | 2639 |
| *zyg-1(it25) ZYG-1<sup>SB+DB3(4A)</sup>* | | 78.9 ± 9.5 | 3373 |
| *zyg-1(it25) fzr-1(bs31)* | | 99.5 ± 1.9 | 1444 |
| *zyg-1(it25)* | 23 | 0.0 ± 0.0 | 880 |
| *zyg-1(it25) ZYG-1<sup>SB(S335A)</sup>* | | 0.8 ± 1.9 | 536 |
| *zyg-1(it25) ZYG-1<sup>SB(T339A)</sup>* | | 1.1 ± 1.9 | 547 |
| *zyg-1(it25) ZYG-1<sup>SB(2A)</sup>* | | 19.6 ± 9.6 | 395 |
| *zyg-1(it25) ZYG-1<sup>DB3(2A)</sup>* | | 20.5 ± 12.5 | 379 |
| *zyg-1(it25) ZYG-1<sup>SB+DB3(4A)</sup>* | | 47.3 ± 26.0 | 566 |
| *zyg-1(it25) fzr-1(bs31)* | | 94.3 ± 12.0 | 718 |
| *zyg-1(it25)* | 24 | 0.0 ± 0.0 | 1010 |
| *zyg-1(it25) ZYG-1<sup>SB(S335A)</sup>* | | 0.0 ± 0.0 | 789 |
| *zyg-1(it25) ZYG-1<sup>SB(T339A)</sup>* | | 0.0 ± 0.0 | 741 |
| *zyg-1(it25) ZYG-1<sup>SB(2A)</sup>* | | 1.6 ± 3.3 | 682 |
| *zyg-1(it25) ZYG-1<sup>DB3(2A)</sup>* | | 1.5 ± 2.4 | 497 |
| *zyg-1(it25) ZYG-1<sup>SB+DB3(4A)</sup>* | | 4.3 ± 3.4 | 627 |
| *zyg-1(it25) fzr-1(bs31)* | | 64.4 ± 15.5 | 830 |
| *zyg-1(it25)* | 25 | 0.0 ± 0.0 | 708 |
| *zyg-1(it25) ZYG-1<sup>SB(2A)</sup>* | | 0.0 ± 0.0 | 627 |
| *zyg-1(it25) ZYG-1<sup>DB3(2A)</sup>* | | 0.0 ± 0.0 | 747 |
| *zyg-1(it25) ZYG-1<sup>SB+DB3(4A)</sup>* | | 0.0 ± 0.0 | 685 |
| *zyg-1(it25) fzr-1(bs31)* | | 2.4 ± 1.7 | 645 |

### Mutating ZYG-1 D-box3 Leads to the *zyg-1* Suppression and Elevated Centrosomal ZYG-1

Next, we utilized the hypomorphic *zyg-1(it25)* background to test how D-box mutation affected ZYG-1 function. In *zyg-1(it25)* mutants, centrosome duplication fails during the first cell cycle, leading to monopolar spindles in second mitosis and 100% embryonic lethality at the restrictive temperature 24°C (O’Connell et al., 2001). If APC/C^FZR-1^ targets ZYG-1 through D-box, mutating functional D-box degron should inhibit APC/C^FZR-1^ binding, thereby protecting ZYG-1 from destruction. Then, ZYG-1 accumulation should compensate for impaired ZYG-1 function in *zyg-1(it25)* mutants. By introducing the same D-box mutations in *zyg-1(it25)* mutants, we asked if any D-box mutation rescued *zyg-1(it25)* phenotypes (**Fig. 2B,C, Table 1**). Intriguingly, the ZYG-1^**DB3(2A)**^ mutation leads to significant restoration of embryonic viability and bipolar spindles to *zyg-1* mutants by >6-fold. Notably, the ZYG-1^**DB3(2A)**^ mutation restores embryonic viability more robustly at the semi-restrictive condition (22.5°C) compared to the restrictive temperature (24°C), suggesting that the ZYG-1^**DB3(2A)**^ mutation requires ZYG-1 activity for the *zyg-1* suppression (**Table 1**).

However, the other D-box mutations increase embryonic lethality and monopolar spindles to *zyg-1(it25)* mutants. It appears that the *zyg-1(it25)* mutation provides a sensitized genetic environment for ZYG-1^**DB1(1A)**^, ZYG-1^**DB2(2A)**^ and ZYG-1^**DB4(1A)**^ mutations as the same ZYG-1 D-box mutations in WT background have little effect on embryonic survival. ZYG-1 function might be impaired by alanine-substitutions at these sites, implicating that these residues are important for ZYG-1 function, presumably as part of the functional domain (**Figs 2A, S2**; Lettman et al., 2013; O’Connell et al., 2001; Shimanovskaya et al., 2014). Although we cannot exclude the possibility of the other D-boxes functioning as degrons, our data support a model where the ZYG-1 D-box3 is a functional degron that mediates the interaction of APC/C coactivator FZR-1 with ZYG-1 for proteasomal degradation.

The ZYG-1^**DB3(2A)**^ mutation should inhibit APC/C^FZR-1^-mediated proteolysis of ZYG-1, leading to ZYG-1 hyperstabilization. To test this, we examined centrosome-associated ZYG-1 by quantitative IF (**Fig. 2D**). Consistent with the *zyg-1* suppression, the ZYG-1^**DB3(2A)**^ mutation leads to elevated centrosomal ZYG-1 levels (1.48 ± 0.4 fold) compared to *zyg-1(it25)* controls (1.00 ± 0.32 fold) while the ZYG-1^**DB2(2A)**^ mutation results in modestly decreased centrosomal ZYG-1 (0.87 ± 0.28 fold), suggesting that the ZYG-1^**DB3(2A)**^ mutation renders ZYG-1 resistant to degradation. It remains unclear whether APC/C^FZR-1^ regulates cellular ZYG-1 levels or affects centrosomal ZYG-1 locally. However, we favor a model where elevated centrosomal ZYG-1 is a direct consequence of increased cellular ZYG-1 levels (Decker et al., 2011; Song et al., 2008). Such correlation between cellular and centrosomal levels of Plk4 has been observed (Guderian et al., 2010).

Since the D-box (1A) mutations may not completely disrupt APC/C^FZR-1^ binding, it remains possible that D-box1 and 4 motifs are functional degrons. To address this, we mutated L352 of D-box3 (349RxxL352) to alanine (ZYG-1^**DB3(1A)**^; **Fig. 2A**), and examined its consequences (**Fig. S1C,D**). The ZYG-1^**DB3(1A)**^ mutation produces neither change on centrosomal ZYG-1 nor rescue of *zyg-1(it25)* lethality, indicating that both R and L residues of D-box3 are required for ZYG-1 destruction, unlike human Cyclin B1 (Matsusaka et al., 2014). Furthermore, the ZYG-1^**DB1(1A)**^ or ZYG-1^**DB4(1A)**^ mutation leads to decreased centrosomal ZYG-1, suggesting alanine-substitutions at these sites negatively impacted ZYG-1 localization (**Fig. S1C,D**). While we cannot exclude the role of D-box1 and 4 as functional degrons, our results support a model where APC/C^FZR-1^ targets ZYG-1 for proteasomal destruction, at least partially through D-box3.

### ZYG-1^DB3(2A)^ Mutation Influences Downstream Effectors of ZYG-1

Increased centrosomal ZYG-1 by the ZYG-1^**DB3(2A)**^ mutation closely correlates with the *zyg-1* suppression. The *zyg-1(it25)* embryo exhibits drastically reduced levels of centrosomal ZYG-1, which negatively affects recruitment of downstream centrosome factors. ZYG-1 is required for SAS-5 and SAS-6 loading to centrosomes, SAS-5 and SAS-6 together recruit SAS-4 (Delattre et al., 2006; Pelletier et al., 2006). SAS-7 was recently identified as centrosome factor critical for centrosomal targeting of SPD-2, and both SAS-7 and SPD-2 act upstream of ZYG-1 (Sugioka et al., 2017).

Elevated centrosomal ZYG-1 by the ZYG-1^**DB3(2A)**^ mutation should restore recruitment of downstream factors, rendering centrosomes competent to assemble new centrioles. To address this, we quantified centrosome-associated protein levels in D-box mutants. To facilitate quantification, we generated epitope-tagged SAS-5::V5 and Ollas::SAS-6 strains at the endogenous locus using CRISPR/Cas9 genome editing (**Fig. S1F,G**). By co-staining embryos with anti-V5 and anti-SAS-4, we first quantified fluorescence intensity of centrosomal SAS-5::V5 and SAS-4 signals (**Fig. 2E**). At anaphase, the ZYG-1^**DB3(2A)**^ mutation leads to elevated centrosomal SAS-5 (1.26 ± 0.39 fold) compared to WT controls (1.00 ± 0.27), which is nearly equivalent to the increase in SAS-5^**KEN(3A)**^ mutants (1.26 ± 0.37 fold) where APC/C^FZR-1^-mediated degradation of SAS-5 is blocked (Medley et al., 2017a). However, centrosomal SAS-5 levels are decreased in ZYG-1^**DB2(2A)**^ mutants (0.70 ± 0.19 fold). Similarly, centrosomal SAS-4 levels are increased in ZYG-1^**DB3(2A)**^ (1.18 ± 0.22 fold) and SAS-5^**KEN(3A)**^ (1.14 ± 0.17 fold) mutants, but reduced in the ZYG-1^**DB2(2A)**^ mutant (0.79 ± 0.22 fold). Furthermore, D-box mutations produce similar effects on SAS-5 and SAS-4 levels in *zyg-1(it25)* mutants **(Fig. S3A,C).** However, SAS-5 and SAS-4 levels are unaffected at metaphase since APC/C^FZR-1^ is active in late mitosis (**Fig. S3B,D).**

Next, we examined SAS-6 levels in ZYG-1 D-box(2A) mutants (**Fig. 2F).** At anaphase, the ZYG-1^**DB3(2A)**^ mutation leads to increased centrosomal SAS-6 (1.26 ± 0.23 fold) compared to WT controls (1.00 ± 0.19 fold), comparable to the change in SAS-5^**KEN(3A)**^ mutants (1.22 ± 0.23 fold), whereas the ZYG-1^**DB2(2A)**^ mutation has little effect on SAS-6 (0.94 ± 0.13 fold). Finally, we monitored dynamic changes of centrosomal GFP::SAS-7 by live imaging (**Fig. S3E**). As expected for an upstream factor, centrosomal SAS-7 levels are unaffected in ZYG-1 D-box mutants. Together, our data show the ZYG-1^**DB3(2A)**^ mutation leads to elevated centrosomal ZYG-1, which promotes centrosomal loading of downstream factors, thereby restoring centrosome duplication and viability to *zyg-1* mutants.

### ZYG-1^DB3(2A)^ Mutation Produces Less Robust Impacts on Centrosomal ZYG-1 and the *zyg-1* Suppression Than Loss of FZR-1

While the ZYG-1^**DB3(2A)**^ mutation leads to elevated centrosomal ZYG-1, it is evident that the *fzr-1* mutation produces greater impact on ZYG-1 levels (1.86 ± 0.44 fold) than the ZYG-1^**DB3(2A)**^ mutation (**Fig. 2D**). Similar trends between *fzr-1(bs31)* and ZYG-1^**DB3(2A)**^ mutations are observed for the *zyg-1* suppression (**Fig. 2B,C, Table 1**).

Since SAS-5 is an APC/C^FZR-1^ substrate and mutating the SAS-5 KEN-box leads to the *zyg-1* suppression (Medley et al., 2017a), we asked if mutating the ZYG-1 D-box3 and SAS-5 KEN-box degrons simultaneously produces the effects comparable to those resulting from the *fzr-1* mutation. Co-mutating ZYG-1 and SAS-5 degrons leads to the *zyg-1* suppression at a level nearly equivalent to the ZYG-1^**DB3(2A)**^ mutation alone (**Fig. 2B,C, Table 1**). Consistently, the effects on centrosomal ZYG-1 levels are comparable between the ZYG-1^**DB3(2A)**^; SAS-5^**KEN(3A)**^ double mutation (1.55 ± 0.42 fold) and ZYG-1^**DB3(2A)**^ single mutation (1.48 ± 0.42 fold; **Fig. 2D**). Since SAS-5 acts downstream of ZYG-1, mutating SAS-5 KEN-box does not affect centrosomal ZYG-1 (1.02 ± 0.3 fold) albeit it increases SAS-5 levels (**Fig. 2D,E**). Thus, it seems unlikely that SAS-5 stabilization had an indirect effect on increased centrosomal ZYG-1.

Our data illustrate the epistatic relationship between *zyg-1* and *sas-5* in centrosome assembly. Co-mutating ZYG-1 and SAS-5 degrons produces the impacts that are comparable to the ZYG-1^**DB3(2A)**^ single mutation and less potent than the *fzr-1* mutation. These results suggest that APC/C^FZR-1^ targets additional centrosome factors, likely acting upstream of ZYG-1, that support ZYG-1 activity. An intriguing candidate is SPD-2 that promotes centrosomal targeting of ZYG-1 (Shimanovskaya et al., 2014). In support of this, *C. elegans* SPD-2 contains putative D-boxes and *Drosophila* Spd2 has been shown to be an APC/C^Fzr/FZR-1^ substrate (Meghini et al., 2016).

### The Conserved Slimb-binding Motif is Critical for Controlling ZYG-1 Levels

While our data suggest APC/C^FZR-1^ controls ZYG-1 stability via D-box3, mutating D-box3 alone produces a modest effect on ZYG-1 levels (**Fig. 2D**), raising the possibility that ZYG-1 levels are regulated via additional degrons (D-boxes or non-canonical degrons) recognized by APC/C, or by other proteolytic pathways. Indeed, ZYG-1 levels are shown to be regulated by SCF^Slimb/βTrCP^-mediated proteolysis (Peel et al., 2012), the mechanism of which is conserved (Cunha-Ferreira et al., 2009; Holland et al., 2010; Rogers et al., 2009). iIn *Drosophila* and human cells, the SCF^Slimb/βTrCP^ E3 ubiquitin ligase controls Plk4 abundance through the conserved Slimb/ßTrCP-binding motif (D**<u>S</u>**Gxx**<u>S</u>/<u>T</u>**) within Plk4, and autophosphorylation of this motif is critical for SCF^Slimb/βTrCP^ binding and Plk4 degradation (Cunha-Ferreira et al., 2013; Guderian et al., 2010; Klebba et al., 2013).

In *C. elegans*, while inhibiting LIN-23 (Slimb/βTrCP homolog) stabilizes ZYG-1, a direct involvement of the ZYG-1 Slimb/ßTrCP-binding (SB) motif (334D**<u>S</u>**Gxx**<u>T</u>**339) in ZYG-1 stability remains unclear (Peel et al., 2012). We thus asked whether the ZYG-1 SB motif influences ZYG-1 levels. Toward this, we generated *zyg-1(it25)* mutant strains carrying alanine-substitution mutations for two critical residues within the ZYG-1 SB motif by CRISPR/Cas9 genome-editing (**Fig. 3A**), equivalent to the known phosphosites (D**S**Gxx**T**) of the SB motif in Plk4 (Cunha-Ferreira et al., 2013; Guderian et al., 2010; Holland et al., 2010; Klebba et al., 2013). Single-alanine substitution (S335A or T339A) leads to ~3-fold increased embryonic viability in *zyg-1(it25)* mutants compared to controls at 22.5°C (**Fig. 3B, Table 1**). Intriguingly, mutating both residues (334D**<u>A</u>**Gxx**<u>A</u>**339), termed the ZYG-1^**SB(2A)**^ mutation, restores embryonic viability to *zyg-1(it25)* mutants by >10-fold higher than controls (**Fig. 3B, Table 1**). Therefore, both S335 and T339 in the ZYG-1 SB motif are critical for ZYG-1 degradation, supporting a model that SCF^Slimb/βTrCP^ targets ZYG-1 for proteasomal degradation through the ZYG-1 SB motif. Our data substantiate the conserved mechanism by which Plk4/ZYG-1 levels are regulated by SCF^Slimb/βTrCP^-mediated proteolysis through the SB motif.

**Figure 3.**
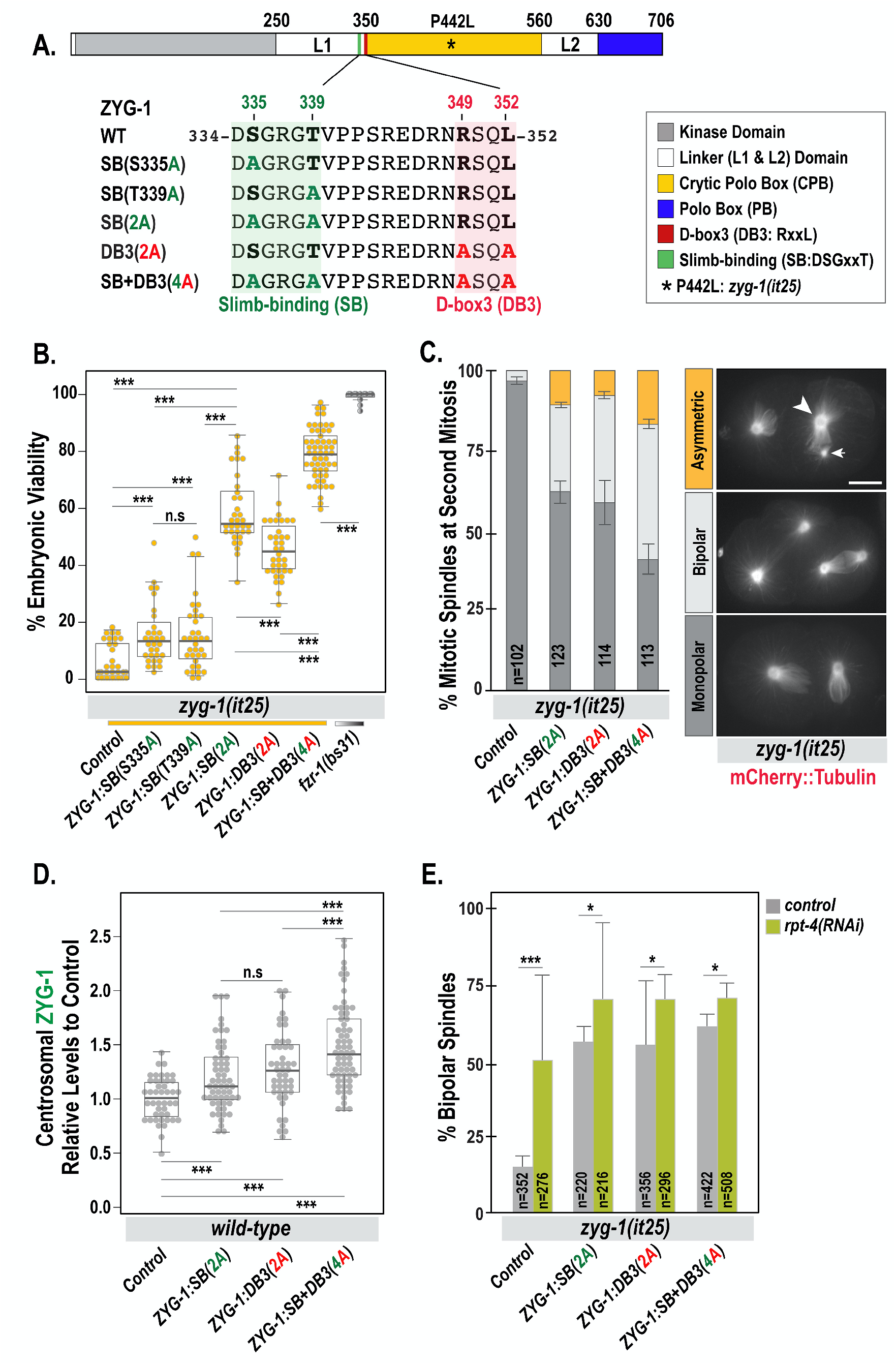
APC/C^FZR-1^ and SCF^Slimb/βTrCP^ regulate ZYG-1 levels cooperatively. (A) ZYG-1 protein structure illustrates the locations of D-box and Slimb-binding motifs and alanine-substitutions. (B) Embryonic viability at 22.5°C (Table 1). Each dot represents a hermaphrodite. (C) Quantification of monopolar (dark grey), bipolar (light grey) and asymmetric (yellow) spindles in two-cell embryos expressing mCherry::Tubulin at 24°C. Arrow and arrowhead indicate asymmetric centrosomes. Bar, 10 μm. (D) Quantification of centrosomal ZYG-1 levels at first mitotic anaphase. Each dot represents a centrosome. (B,D) In the plots, box ranges from the first through third quartile of the data. Thick bar indicates the median. Solid grey line extends 1.5 times the inter-quartile range or to the minimum and maximum data point. (E) Quantification of bipolar spindles in two-cell embryos at 24°C. (C,E) n is the number of blastomeres. Average and SD are presented. (B,D,E) ^ns^*p*>0.05, \**p*<0.05, \*\*\**p*<0.001 (two-tailed t-tests).

### APC/C^FZR-1^ and SCF^Slimb/βTrCP^ E3 Ubiquitin Ligases Regulate ZYG-1 Levels Cooperatively

Given that both APC/C^FZR-1^ and SCF^Slimb/βTrCP^ regulate ZYG-1 levels, we asked if co-mutating ZYG-1 SB and D-box3 motifs could enhance ZYG-1 protection from degradation, and produced the strain carrying double-mutation of ZYG-1 SB and D-box3 motifs at the endogenous locus (ZYG-1^**SB+DB3(4A)**^; **Fig. 3A**). The ZYG-1 SB (334DSGxxT339) and D-box3 (349RxxL352) motifs reside in close proximity within the L1 domain (**Fig. 3A**). Whereas neither motif is conserved in closely related nematodes (**Fig. S2D**), the flanking region of these motifs appears to be somewhat conserved (Fig. 2D in Klebba et al., 2013).

In *zyg-1(it25)* background, mutating both ZYG-1 SB and D-box3 motifs further increases embryonic viability and bipolar spindles, compared to ZYG-1^**SB(2A)**^ or ZYG-1^**DB3(2A)**^ single mutation (**Fig. 3B,C, Table 1**). Consistently, mutating both degrons leads to further elevated centrosomal ZYG-1 levels (1.52 ± 0.37 fold) than ZYG-1^**SB(2A)**^ (1.20 ± 0.31 fold) or ZYG-1^**DB3(2A)**^ (1.29 ± 0.34 fold) mutation (**Fig. 3D**). Thus, inhibiting both APC/C^FZR-1^ and SCF^Slimb/βTrCP^ ubiquitin ligases enhances ZYG-1 stability in an additive manner, suggesting that APC/C^FZR-1^ and SCF^Slimb/βTrCP^-mediated proteolytic regulation cooperatively controls ZYG-1 stability via conserved motifs.

Finally, we asked if ZYG-1^**DB3(2A)**^ mutation influences ZYG-1 function through the 26S proteasome (**Fig. 3E**). We depleted the proteasome component RPT-4 using *rpt-4(RNAi)* to partially inhibit proteasome function, which still allows completion of meiosis and mitotic cycles (Song et al., 2011). Given the observation that *rpt-4(RNAi)* leads to increased centrosomal ZYG-1 (Peel et al., 2012), we examined how *rpt-4(RNAi)-mediated* inhibition of the proteasome affected ZYG-1 function in centrosome duplication. As expected, *rpt-4(RNAi)* in *zyg-1(it25)* mutants significantly restores bipolar spindles compared to control RNAi (50 ± 28% vs 15 ± 4%; 3.33 fold). By contrast, *rpt-4(RNAi)* combined with ZYG-1^**DB3(2A)**^ mutation in *zyg-1(it25)* mutants leads to a small increase in bipolarity relative to control RNAi (70 ± 8% vs 55 ± 21%; 1.16 fold). Similarly, *rpt-4(RNAi)* with ZYG-1^**SB(2A)**^ (70 ± 25% vs 56 ± 5%; 1.25 fold) or ZYG-1^**SB+DB3(4A)**^ (70 ± 5% vs 61 ± 4%; 1.15 fold) mutation exhibits slightly increased bipolarity compared to controls. These results further support that ZYG-1^DB3(2A)^ mutation renders ZYG-1 partially resistant to proteasomal degradation, consistent with ZYG-1^**SB(2A)**^ and ZYG-1^**DB3(2A)**^ mutations partially blocking proteolysis, leading to ZYG-1 stabilization and elevated centrosomal ZYG-1.

In summary, our work reports, for the first time in any organism, that centrosomal ZYG-1/Plk4 levels are influenced by APC/C^FZR-1^. Our data also support that two E3 ubiquitin ligases, APC/C^FZR-1^ and SCF^Slimb/βTrCP^, cooperate to facilitate timely degradation of ZYG-1 during the rapid cell cycle in *C. elegans* embryos, preventing aberrant centrosome numbers (**Fig. S1H,I**). Along with proteolytic regulation of ZYG-1 levels, ZYG-1 activity is shown to be modulated by kinase and phosphatases (Kitagawa et al., 2011; Medley et al., 2017b; Peel et al., 2017; Song et al., 2011). Thus, multiple mechanisms appear to be involved in precise control of ZYG-1 activity, which might account for rare observations of centrosome amplification in our ZYG-1 degron mutants (**Fig. S1I**). Interestingly, centrosome amplification phenotypes were observed in neither *fzr-1* nor SAS-5^**KEN(3A)**^ mutants (Medley et al., 2017a). Finally, our results suggest that APC/C^FZR-1^ targets additional centrosome factors acting upstream or parallel of ZYG-1. Additional work is required to further understand the complete regulatory mechanisms of APC/C^FZR-1^ in centrosome assembly. It will be also interesting to explore the role of APC/C^cdh1/FZR-1^ in regulating Plk4 stability and its association with human cancers.

## Materials and Methods

### *C. elegans* Culture and Genetic Analysis

The *C. elegans* strains used in this study are listed in Table S1. All strains were derived from the wild-type Bristol N2 strain (Brenner, 1974, Church et al., 1995) and maintained on MYOB plates seeded with *Escherichia coli* OP50 at 16 or 19°C. Some strains were provided by the CGC, which is funded by NIH Office of Research Infrastructure Programs (P40 OD010440). For genetic analysis, individual L4 hermaphrodites were transferred to new plates and allowed to produce progeny for 24-48 h at the temperatures indicated. Progeny were allowed to develop for 18-24 h before counting the number of larvae and dead eggs. RNAi feeding was performed as described (Kamath et al., 2003) and the L4440 empty feeding vector was used as a negative control. Animals were fed RNAi for 12-18h prior to imaging.

### Immunofluorescence and Cytological analysis

Immunofluorescence and confocal microscopy were performed as described (Song et al., 2008). The following primary and secondary antibodies were used at 1:3000 dilutions: DM1a (Sigma, #T9026), α-ZYG-1 (Stubenvoll et al., 2016), α-SAS-4 (Song et al., 2008), α-Myc (ThermoFisher, #PA1-981; **Fig. S1E**), and α-V5 (MBL, #M167-3; **Fig. S1F**), α-Ollas (ThermoFisher, #MA5-16125; **Fig. S1G**), Alexa Fluor 488 and 568 secondary antibodies (ThermoFisher, #A11001, A11004, A11006, A11034, A11036). Time-lapse recordings for GFP::SAS-7 embryos were performed as described previously (Medley et al. 2017b). Images of GFP::SAS-7 were acquired at 30 second intervals starting from prometaphase until the completion of first cell division. Confocal microscopy was performed using a Nikon Eclipse Ti-U microscope equipped with a Plan Apo 60×1.4 NA lens, a Spinning Disk Confocal (CSU X1) and a Photometrics Evolve 512 camera. MetaMorph software (Molecular Devices, Sunnyvale, CA, USA) was used for image acquisition and quantification of the fluorescence intensity, and Adobe Photoshop/Illustrator 2020 for image processing. To quantify centrosomal signals, the average intensity within 8 or 9-pixel (1 pixel=0.151 μm) diameter region was recorded for the highest intensity of the focal plane within an area centered on the centrosome. The average intensity within a 25-pixel diameter region drawn outside of the embryo was used for background subtraction.

### CRISPR/Cas9 Genome Editing

For genome editing, we used the co-CRISPR technique described previously (Arribere et al., 2014, Paix et al., 2015). crRNA was designed using the CRISPOR web server (crispor.tefor.net; Concordet and Haeussler, 2018). Animals were microinjected with a mixture of commercially available SpCas9 (IDT, Coralville, IA) and custom-designed oligonucleotides (**Tables S2,S3**) including crRNAs at 0.4–0.8 μg/μl, tracrRNA at 12 μg/μl, and single-stranded DNA oligonucleotides at 25–100 ng/μl. The amount of crRNA was tripled for low-efficiency crRNAs (ZYG-1 D-box1 and D-box4, Ollas::FZR-1). After injection, we screened for *dpy-10(cn64) II/+* rollers in F1 progeny and genotyping F2. The genome editing was verified by Sanger Sequencing (GeneWiz, South Plainfield, NJ).

### Immunoprecipitation (IP) and Immunoblot

IP experiments were performed as described in (Stubenvoll et al., 2016). Embryos were collected from young gravid worms using hypochlorite treatment (1:2:1 ratio of M9 buffer, 5.25% sodium hypochlorite, and 5 M NaOH), frozen in liquid nitrogen and stored at −80°C until use. Embryos were suspended in lysis buffer [50 mM HEPES, pH 7.4, 1 mM EDTA, 1 mM MgCl2, 200 mM KCl, and 10% glycerol (v/v)] with complete protease inhibitor cocktail (Roche) and MG132 (Tocris, Avonmouth, Bristol, UK), milled for 5 minutes (repeat x3) at 30 Hz using a Retsch MM 400 mixer-mill (Verder Scientific, Newtown, PA), then sonicated for 3 minutes in ultrasonic bath (Thermo-Fisher). Lysates were spun at 45,000 rpm for 45 minutes using a Sorvall RC M120EX ultracentrifuge (Thermo-Fisher), then the supernatant was recovered to clean tubes. The equivalent amount of total protein lysates was used for IP. Embryonic protein lysates mixed with α-GFP or α-Myc magnetic beads (MBL, # D153-11; M047-11) were incubated by rotation for 1 hour at 4°C, and washed (3x 5 minutes) with PBST (PBS + 0.1% Triton-X 100). IP with beads and input samples were resuspended in 2X Laemmli Sample Buffer (Sigma), and boiled for 5 minutes before fractionating on a 4–12% NuPAGE Bis-Tris gel (Invitrogen). Proteins on a gel were transferred to a nitrocellulose membrane and analyzed by using the antibodies at 1:3000-10,000 dilutions: DM1a (Sigma, #T9026), α-GFP (Roche, #11814460001), α-Myc (Genscript, #A00704), α-SAS-5 (Medley et al., 2017), IRDye secondary antibodies (LI-COR Biosciences). Blots were imaged using the Odyssey infrared scanner (LI-COR Biosciences), and analyzed using Image Studio software (LI-COR Biosciences).

### Statistical analysis

Statistics were produced using R statistical software and presented as average ± standard deviation (SD). Dotplots were generated using the R ‘beeswarm’ package. In the dotplots, box ranges from the first through third quartile of the data. Thick bar indicates the median. Solid grey line extends 1.5 times the inter-quartile range or to the minimum and maximum data point. All P-values were calculated using two-tailed t-tests: ^ns^*p*>0.05, \**p*<0.05, \*\**p*<0.01, \*\*\**p*<0.001.

