## Supplemental Information for "APC/C^FZR-1^ Controls ZYG-1 Levels to Regulate Centrosome Assembly"

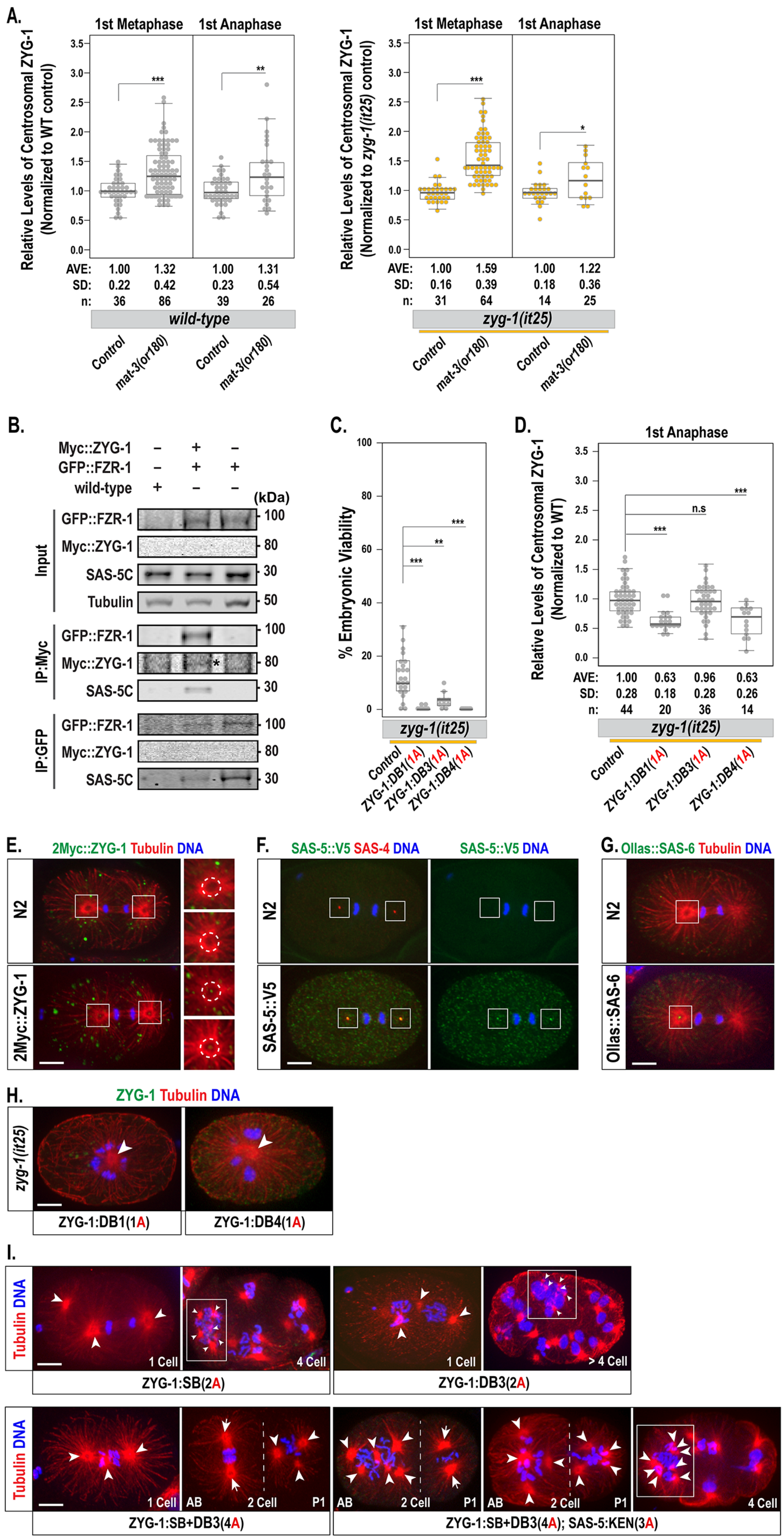

A.
1st Metaphase
1st Anaphase
Relative Levels of Centrosomal ZYG-1 (Normalized to WT control)
AVE: 1.00 1.32 1.00 1.31
SD: 0.22 0.42 0.23 0.54
n: 36 86 39 26
wild-type
Control
mat-3(or180)
Relative Levels of Centrosomal ZYG-1 (Normalized to zyg-1(it25) control)
AVE: 1.00 1.59 1.00 1.22
SD: 0.16 0.39 0.18 0.36
n: 31 64 14 25
zyg-1(it25)
B.
Myc::ZYG-1
GFP::FZR-1
wild-type
(kDa)
Input
IP:Myc
IP:GFP
GFP::FZR-1
Myc::ZYG-1
SAS-5C
Tubulin
100
80
30
50
C.
% Embryonic Viability
zyg-1(it25)
Control
ZYG-1:DB1(1A)
ZYG-1:DB3(1A)
ZYG-1:DB4(1A)
D.
1st Anaphase
Relative Levels of Centrosomal ZYG-1 (Normalized to WT)
n.s
AVE: 1.00 0.63 0.96 0.63
SD: 0.28 0.18 0.28 0.26
n: 44 20 36 14
zyg-1(it25)
E.
2Myc::ZYG-1 Tubulin DNA
N2
2Myc::ZYG-1
F.
SAS-5::V5 SAS-4 DNA
SAS-5::V5 DNA
N2
SAS-5::V5
G.
Ollas::SAS-6 Tubulin DNA
N2
Ollas::SAS-6
H.
ZYG-1 Tubulin DNA
zyg-1(it25)
ZYG-1:DB1(1A)
ZYG-1:DB4(1A)
I.
Tubulin DNA
1 Cell
4 Cell
> 4 Cell
ZYG-1:SB(2A)
ZYG-1:DB3(2A)
AB
2 Cell
P1
ZYG-1:SB+DB3(4A)
ZYG-1:SB+DB3(4A); SAS-5:KEN(3A)

**Fig S1**. (**A**) Quantification of centrosomal ZYG-1 levels in wild-type and *mat-3(or180)* mutants at 21°C (left), and *zyg-1(it25)* and *zyg-1(it25); mat-3(or180)* mutants at 22.5°C (right). (**B**) Immunoprecipitation (IP) data suggest that FZR-1 physically interacts with ZYG-1. GFP::FZR-1 and 2xMyc::ZYG-1(*) are detected in the 2xMyc::ZYG-1 pulldown materials. SAS-5 is also detected in the 2xMyc::ZYG-1/GFP::FZR-1 complex, consistent with the observation that SAS-5 and FZR-1 physically interact (Medley et al., 2017). ~10 % of total embryonic lysates was loaded in the input lanes. Tubulin is used as loading control for input samples. (**C-D**) Both R and L residues in D-box3 motif (RxxL) are critical for ZYG-1 degradation. (**C**) % Embryonic viability of the *zyg-1(it25)* mutants carrying single-alanine substitutions of the ZYG-1 D-box (Rxx**A**) animals grown at 22.5°C. Each dot represents a hermaphrodite. (**D**) Quantification of centrosomal ZYG-1 levels in the *zyg-1(it25)* mutants carrying single-alanine substitutions of the ZYG-1 D-box (1A: Rxx**A**) at 23°C. (**A,D**) Each dot represents a centrosome. (**A,C,D**) Box ranges from the first through third quartile of the data. Thick bar indicates the median. Solid grey line extends 1.5 times the inter-quartile range or to the minimum and maximum data point. $^{ns}p>0.05$, $^{*}p<0.05$, $^{**}p<0.01$, $^{***}p<0.001$ (two-tailed t-test). (**E-G**) The specificity of antibodies: The embryos expressing N-terminal 2xMyc tagged ZYG-1 (**E**), a C-terminal V5 tagged SAS-5 (**F**) or N-terminal Olla-tagged SAS-6 (**G**) at endogenous levels from the native genomic locus generated using the CRISPR/Cas-9 method (see **Table S2,S3**): $\alpha$-Tubulin (**E,G**) and SAS-4 (**F**) as centrosome marker. Antibodies against Myc, V5 and Ollas-tag detect 2xMyc::ZYG-1 (**E**), SAS-5::V5 (**F**) and Ollas::SAS-6 (**G**) at centrioles, but no centriolar signals are detected in wild-type (N2) embryos (negative control). (**H,I**) Abnormal numbers of centrosomes are observed in the ZYG-1 degron mutants: (**H**) Mutating the ZYG-1 D-box1 and D-box4 in *zyg-1(it25)* mutants leads to more frequent monopolar spindles in one-cell embryos than the *zyg-1(it25)* controls grown at semi-restrictive temperature conditions (<24°C). (**I**) Mutating ZYG-1 Slimb-binding (SB) and/or D-box3 (DB3), or combined with the SAS-5 KEN-box mutation results in extra centrosomes (>2 centrosomes) during early embryonic cell division albeit at extremely rare occasions (3-5%, n >500) in the ZYG-1 degron mutants tested here. Arrowheads indicate abnormal centrosomes and arrows illustrate bipolar spindles: Dashed line indicates cell boundary. In 4-cell or > 4-cell stage embryos, a cell with extra centrosomes is box-highlighted. (**E-I**) Bar, 10 $\mu$m.

A. Protein Structure of ZYG-1 Related Proteins

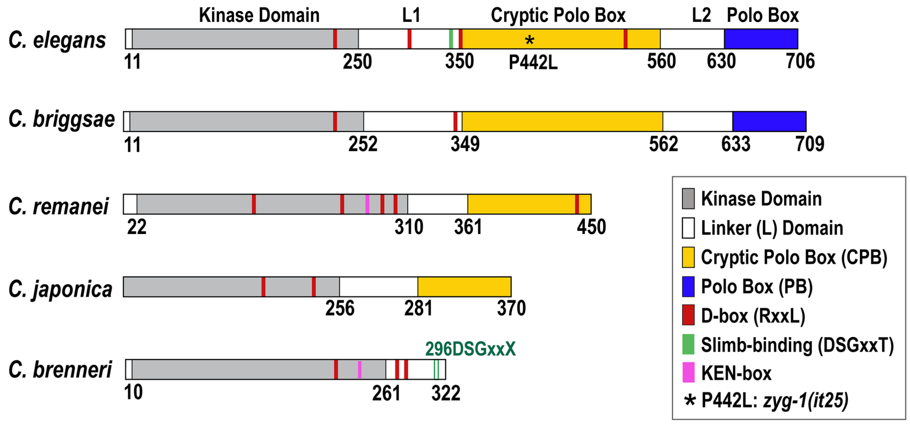


B. ZYG-1 D-box1 (237RIPL240)

```
                            D-box1
C. elegans   227 EQMMDTDAKKRIPLTQIVLSEFMYENTNENA 257
C. briggsae  229 ERMMHTDARRRIQLKEIVMTDYVKAKMGEAT 259
C. remanei   240 KRMMETNPKRRIELREIVMSEFMKENMDEEG 270
C. japonica  168 EAMMHTDAVRRIKLKEIVMTDFMRENTDDEA 198
C. brenneri  227 EKMMHANVKERIPLREIVMTDYMKENTDDEA 257
```

C. ZYG-1 D-box2 (303RAPL306)

```
                 Positively Charged Patch (SAS-6 binding: Lettman et al., 2013)                  D-box2
C. elegans   260 FSREHSRDGRRQRSREPVRSSRDDRSRDGRALIRSSSQPAHSGRAPL 306
C. briggsae  261 GSREHSRDSRSQRSREPFRSSRDGISLERRPPARSSSQPVNSRRDPD 307
C. remanei   272 FSREHSKDSRHQLSREPRISSRDERSQDRR-PLRSSSQPVNSARMTH 317
C. japonica  201 YSRESSRDGRR-RSREPRYPSRDGRSQDRRPPLRSSSQPVNSARMPM 246
C. brenneri  261 YSREHSRDSRQ-RSREPIR-SRDALSQDRKPLTRYPEKRQAHLILYC 305
```

D. ZYG-1 D-box3 (349RSQL352)

```
                           Slimb-binding (334..339)  D-box3
C. elegans   323 FDSERGRERDRDSGRGTVPPSREDRNRSQLWPIRMD 358
C. briggsae  323 VEPDRARVRHRLSARG-IGSSQEDDLRQQIWPIRME 357
C. remanei   335 SENDRGRARQRTSARG-LGTSHETQSREQIWPIRME 369
C. japonica  267 MPT-------------PLPSGERDRHHNNMWPIRME 289
```

E. ZYG-1 D-box4 (489RIKL492) Adapted from Shimanovskaya et al., 2013

```
                              D-box4
C. elegans   477 DAQAQLMENGDLRIKLPRSVIV 498
C. briggsae  478 DASAQLMENGDLRIRFPNS-LI 498
B. malayi    555 DSTARLMENGDFRIKFQDGRLA 576
L. loa       534 DSTARLMENGDFRVKFQDGRLA 555
A. suum      551 ETTARLMENGDFRLKFSDGRLA 572
```

**Fig S2. Sequence Alignments of ZYG-1 Related Proteins in Nematodes** (A) The closely related *Caenorhabditis* ZYG-1 protein structures illustrates the functional domains and the locations of putative degron motifs (D-box, KEN-box and Slimb-binding motifs) (O'Connell et al., 2001, Lettman et al., 2013, Shimanovskaya et al., 2014, Peel et al., 2012). (B-D) Sequence alignments of putative D-box flanking regions in ZYG-1 related proteins. Amino acid residues in red indicate critical sites for D-box motifs (**R**xx**L**). Sites in green highlight the ZYG-1 Slimb-binding degron motif (D**S**Gxx**T**). (C) The *C. elegans* ZYG-1 D-box2 and SAS-6 binding region (Lettman et al., 2013) are located in close proximity within L1 domain. Positively charged residues are highlighted in bold. (B-E) The WormBase IDs of the aligned sequences: *C. elegans*, CE28571; *C. briggsae*, BP:CBP14405; *C. remanei*, RP:RP38674; *C. brenneri*, CN:CN31126; *C. japonica*, JA:JA58610. Alignments were performed using the Clustal Omega (https://www.ebi.ac.uk/Tools/msa/clustalo/) and COBALT (https://www.ncbi.nlm.nih.gov/tools/cobalt/re_cobalt.cgi) tools. (E) The *C. elegans* D-box4 motif flanking region is aligned in other nematode homologs (adapted from Shimanovskaya et al., 2014): *Brugia malayi; Loa loa; Ascaris suum*.

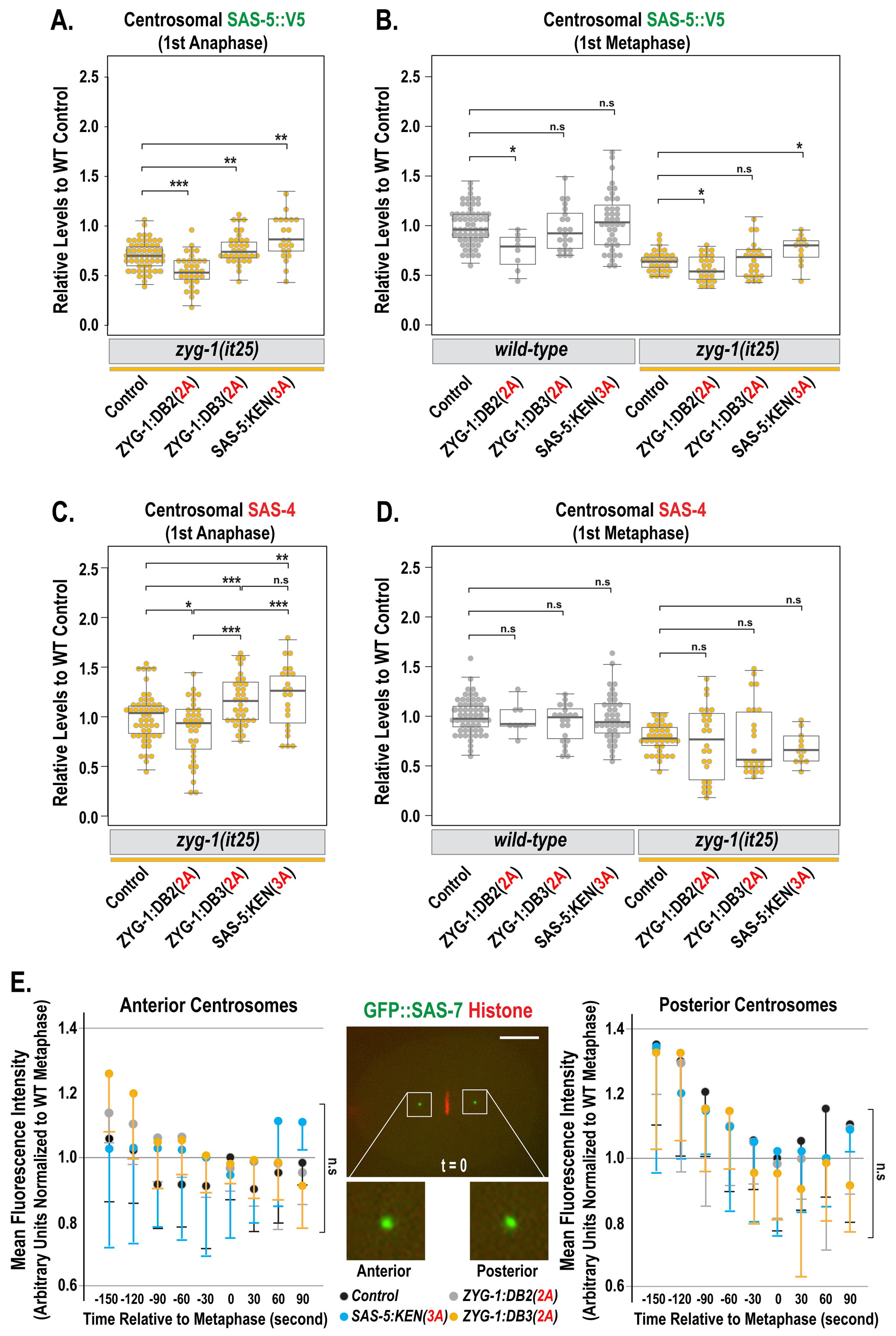


**Fig S3. Quantification of SAS-5 (A-B), SAS-4 (C-D) and SAS-7 (E) levels** at anaphase *zyg-1(it25)* mutant centrosomes (A,C) and at metaphase wild-type centrosomes (B,D) affected by ZYG-1 D-box and SAS-5 KEN-box mutations. (A-D) Each dot represents a centrosome. Box ranges from the first through third quartile of the data. Thick bar indicates the median. Solid grey line extends 1.5 times the inter-quartile range or to the minimum and maximum data point. $^{ns}p>0.05$, $^{*}p<0.05$, $^{**}p<0.01$, $^{***}p<0.001$ (two-tailed t-test). (E) Measurements of fluorescence intensity of GFP::SAS-7 at anterior (left panel) and posterior (right panel) centrosomes during the first cell cycle. Data were collected from the number of centrosomes (n)=5-13 per timepoint for each strain. Mean (dots) - s.d. (vertical lines) are presented relative to the first metaphase (t=0). Shown is the first metaphase embryo expressing GFP::SAS-7 and mCherry::Histone at 24°C (middle panel). Bar, 10 μm.

### Table S1. List of *C. elegans* Strains

| Strain | Genotype | Origin |
|---|---|---|
| **N2** | wild-type | CGC |
| **MTU6** | *fzr-1(bs31) II* | Medley et al., 2017 |
| **OC14** | *zyg-1(it25[ZYG-1*$^{P442L}$*]) II* | O'Connell et al., 2001 |
| **OC201** | *zyg-1(it25) fzr-1(bs31) II* | Kemp et al., 2007 |
| **MTU7** | *mat-3(or180) III* | Golden et al., 2000 |
| **MTU8** | *zyg-1(it25) II; mat-3(or180) III* | Medley et al., 2017 |
| **MTU13** | *zyg-1(it25) II; sas-5(mhs359[SAS-5*$^{KEN(3A)}$*]) V* | Medley et al., 2017 |
| **MTU350** | *zyg-1(mhs584[ZYG-1*$^{DB1(1A)}$*]) II* | This Study |
| **MTU353** | *zyg-1(mhs584it25) II* | This Study |
| **MTU388** | *zyg-1(mhs437[ZYG-1*$^{DB2(2A)}$*]) II* | This Study |
| **MTU367** | *zyg-1(mhs449it25[ZYG-1*$^{DB2(2A):P442L}$*]) II* | This Study |
| **MTU288** | *zyg-1(mhs449it25) II; sas-5(mhs359) V* | This Study |
| **MTU172** | *zyg-1(mhs476[ZYG-1*$^{DB3(2A)}$*]) II* | This Study |
| **MTU439** | *zyg-1(mhs498it25[ZYG-1*$^{DB3(2A):P442L}$*]) II* | This Study |
| **MTU406** | *zyg-1(mhs614it25[ZYG-1*$^{DB3(1A):P442L}$*]) II* | This Study |
| **MTU329** | *zyg-1(mhs498it25) II; sas-5(mhs359) V* | This Study |
| **MTU324** | *zyg-1(mhs575[ZYG-1*$^{DB4(1A)}$*]) II* | This Study |
| **MTU387** | *zyg-1(it25mhs592[ ZYG-1*$^{P442L:DB4(1A)}$*]) II* | This Study |
| **MTU150** | *sas-6(mhs451[Ollas::SAS-6]) IV* | This Study |
| **MTU595** | *zyg-1(mhs437) II; sas-6(mhs451) IV* | This Study |
| **MTU593** | *zyg-1(mhs476) II; sas-6(mhs451) IV* | This Study |
| **MTU594** | *sas-6(mhs451) IV; sas-5(mhs359) V* | This Study |
| **MTU257** | *sas-5(mhs533[SAS-5::V5]) V* | This Study |
| **MTU422** | *zyg-1(mhs437) II; sas-5(mhs533) V* | This Study |
| **MTU423** | *zyg-1(mhs476) II; sas-5(mhs533) V* | This Study |
| **MTU357** | *sas-5(mhs359mhs568[SAS-5*$^{KEN(3A)}$*::V5]) V* | This Study |
| **MTU354** | *zyg-1(it25) II; sas-5(mhs533) V* | This Study |
| **MTU420** | *zyg-1(mhs449it25) II; sas-5(mhs533) V* | This Study |
| **MTU429** | *zyg-1(mhs498it25) II; sas-5(mhs533) V* | This Study |
| **MTU496** | *zyg-1(it25) II; sas-5(mhs359mhs568) V* | This Study |
| **MTU271** | *zyg-1(mhs456[2xMyc::ZYG-1]) II* | This Study |
| **MTU309** | *zyg-1(mhs561mhs524[2xMyc::ZYG-1*$^{SB(2A)}$*]) II* | This Study |
| **MTU419** | *zyg-1(mhs603mhs476[2xMyc::ZYG-1*$^{DB3(2A)}$*]) II* | This Study |
| **MTU553** | *zyg-1(mhs603mhs675mhs476[2xMyc::ZYG-1*$^{SB+DB3(4A)}$*]) II* | This Study |
| **MTU649** | *zyg-1(mhs603mhs675mhs476[2xMyc::ZYG-1*$^{SB+DB3(4A)}$*]) II; sas-5(mhs359[SAS-5*$^{KEN(3A)}$*]) V* | This Study |
| **MTU403** | *zyg-1(mhs629it25[ZYG-1*$^{S335A:P442L}$*]) II* | This Study |
| **MTU445** | *zyg-1(mhs526it25[ZYG-1*$^{T339A:P442L}$*]) II* | This Study |
| **MTU617** | *zyg-1(mhs557it25[ZYG-1*$^{SB(2A)}$*]) II* | This Study |
| **MTU584** | *zyg-1(mhs669mhs498it25[ZYG-1*$^{SB+DB3(4A):P442L}$*]) II* | This Study |
| **EU3000** | *sas-7(or1940[GFP::SAS-7]) III; itls37 IV* | Sugioka et al., 2017 |
| **MTU591** | *zyg-1(mhs437) II; sas-7(or1940) IV* | This Study |
| **MTU599** | *zyg-1(mhs476) II; sas-7(or1940) IV* | This Study |
| **MTU592** | *sas-7(or1940) IV; sas-5(mhs359) V* | This Study |
| **OC190** | *unc-119(ed3) III; bsIs6 [unc-119(+), pie-1p::gfp::fzr-1]* | Medley et al., 2017 |
| **MTU664** | *zyg-1(mhs456[2xMyc::ZYG-1]) II; unc-119(ed3) III; bsIs6 [unc-119(+), pie-1p::gfp::fzr-1]* | This Study |
| **OC481** | *unc-119(ed3) III; bsIs15[pNP99:unc-119(+) tbb-1p::mCherry::tbb-2::tbb-2 3'-utr]* | Medley et al., 2017 |
| **MTU631** | *zyg-1(it25) II; unc-119(ed3) III; bsIs15[pNP99:unc-119(+) tbb-1p::mCherry::tbb-2::tbb-2 3'-utr]* | This Study |
| **MTU476** | *zyg-1(mhs557it25) II; unc-119(ed3) III; bsIs15[pNP99:unc-119(+) tbb-1p::mCherry::tbb-2::tbb-2 3'-utr]* | This Study |
| **MTU573** | *zyg-1(mhs497it25) II; unc-119(ed3) III; bsIs15[pNP99:unc-119(+) tbb-1p::mCherry::tbb-2::tbb-2 3'-utr]* | This Study |
| **MTU563** | *zyg-1(mhs669mhs498it25) II; unc-119(ed3) III; bsIs15[pNP99:unc-119(+) tbb-1p::mCherry::tbb-2::tbb-2 3'-utr]* | This Study |

### Table S2. List of crRNA for CRISPR/Cas9 Genome Editing

| Gene | Target | Sequence (5'-3') |
|---|---|---|
| ***dpy-10*** (Arribere et al., 2014) | *dpy-10(cn64)* co-CRISPR | UUCUGCUGUCUUGAUUGACG |
| ***fzr-1*** | N-terminus | UUACCCUUUUGCACAUAAAA |
| ***sas-5*** | C-terminus | UUCGUGAAAAAUACGCUCGC |
| ***sas-6*** | N-terminus | GCUCAACGGCUGUAGAAGUG |
| ***zyg-1*** | N-terminus | UUGAUCAACUAUGAGAUGAG |
| ***zyg-1*** | D-box1 | GAUAAAACUAUCUGCGUUAA |
| ***zyg-1*** | D-box2 | UGAAUUGGUCGAUUGCUCAG |
| ***zyg-1*** | D-box3 | GGAUCGGCCACAACUGUGAA |
| ***zyg-1*** | D-box4 | CUGAUCUCGGAAGUUUGAUA |
| ***zyg-1*** | Slimb-binding | GGCUUGGAGGUACGGUACCG |

### Table S3. List of ssODN Homologous Repair Templates for CRISPR/Cas9 Genome Editing

| Gene | Variation | Sequence (5'-3') |
|---|---|---|
| ***dpy-10***<br>(Arribere et al., 2014) | *dpy-10(cn64)* | CACTTGAACTTCAATACGGCAAGATGAGAATGACTGGA<br>AACCGTACCGCATGCGGTGCCTATGGTAGCGGAGCTTC<br>ACATGGCTTCAGACCAACAGCCTAT |
| ***fzr-1*** | N-terminal Ollas Tag | GCTTTTGCGTGTTCTCCTCAACTTTACCCTTTTGCACAT<br>AAAATGTCTGGATTCGCTAACGAGCTTGGACCACGTCTT<br>ATGGGAAAGGGAGGTTCCGGTGGTTCTGGTGGATCCG<br>ATGAGCAGCAACCGCCAGCCAACTCTCCGGCTATTTTT<br>CATTCGC |
| ***sas-5*** | C-terminal V5 Tag | GTACCTGAGACTCCAGCTGAACGAGAACGCCGTATTCG<br>TGAAAAATACGCTCGCCGTAAAGGAGGTTCCGGTGGTT<br>CTGGTGGATCCGGTAAGCCTATCCCAAATCCTTTGTTG<br>GGTCTGGACTCCACGTGATATCAAATGTGTTTAACTCTT<br>GACGTTTTAAAATTATCATTAATCACTACG |
| ***sas-6*** | N-terminal Ollas Tag | CTTACTTAAAATCCGGTTGATTTAGGCTCAACGGCTGTA<br>GAAGTGATGCGATGAGTGTTTGATCGAATAATGCAATTT<br>TGCTAGTCTTTCCCATAAGACGTGGTCCAAGCTCGTTAG<br>CGAATCCAGACATTTTTGTGGGAGAAAATCTGAAAAAAA<br>TTATGTTAAAAATAG |
| ***zyg-1*** | N-terminal 2xMyc Tag | TTTTAGCGCACCAAGTGTTGATCAACTATGAGATGGAGC<br>AGAAGTTGATAAGTGAAGAAGACTTAGAGCAGAAATTGA<br>TTTCGGAGGAGGATTTGGGAGGCTCGGGCGGTTCGGG<br>AGGATCTAGCGGTGGGAAGAGTGGTTCAAGATTGAGTG<br>TAGG |
| ***zyg-1*** | D-Box1(AxxA) | GGAACAAATGATGGATACAGATGCAAAGAAGGCTATTC<br>CGGCTACCCAAATTGTCTTATCAGAGTTCATGTACGAGA<br>ATACGAATGAAAATGC |
| ***zyg-1*** | D-Box1(RxxA) | GGAACAAATGATGGATACAGATGCAAAGAAGCGTATTC<br>CGGCTACCCAAATTGTCTTATCAGAGTTCATGTACGAGA<br>ATACGAATGAAAATGC |
| ***zyg-1*** | D-Box2(AxxA) | TGATAAGGTCTTCGAGTCAACCTGCACATTCTGGCGCT<br>GCACCGGCTTCTAACCGCCCAATTCATGACAGGGTAAG<br>TGTTCCAGCAGATCA |
| ***zyg-1*** | D-Box3(AxxA) | GCGGTACCGTACCTCCAAGCCGTGAAGATCGAAACGCT<br>TCACAGGCTTGGCCGATCCGAATGGATAGGTTAGAAGG<br>ACAAAG |
| ***zyg-1*** | D-Box3(RxxA) | GCGGTACCGTACCTCCAAGCCGTGAAGATCGAAACAGA<br>TCACAGGCCTGGCCGATCCGAATGGATAGGTTAGAAGG<br>ACAAAG |
| ***zyg-1*** | D-Box4(AxxA) | GATGCTCAAGCTCAACTTATGGAGAATGGAGATCTCGC<br>TATCAAAGCTCCGAGATCAGTAATTGTTCGTAAAATGGA<br>TAATGG |
| ***zyg-1*** | D-Box4(RxxA) | GATGCTCAAGCTCAACTTATGGAGAATGGAGATCTCAG<br>AATCAAAGCCCCGAGATCAGTAATTGTTCGTAAAATGGA<br>TAATGG |
| ***zyg-1*** | Slimb-binding(S335A) | GGATTTGATTCTGAGAGAGGAAGAGAACGGGATAGAGA<br>CGCCGGACGCGGTACCGTACCTCCAAGCCGTGAAGAT<br>CGAAACCG |
| ***zyg-1*** | Slimb-binding(S335A:T339A) | GAGAGAGGAAGAGAACGGGATAGAGACGCCGGACGCG<br>GTGCTGTACCTCCAAGCCGTGAAGATCGAAACCGTTCA<br>CAG |
